# The sperm-associated microbiota of crickets and their local variation and rapid turnover

**DOI:** 10.1101/2021.10.10.463754

**Authors:** Barbara A. Eckel, Anastasia V. Illner, Oliver Otti, Klaus Reinhardt

## Abstract

While studying aspects of the sperm biology and immunity of two species of crickets, we encountered bacteria that were released from the male sperm container, the spermatophore, alongside sperm. We describe a presumably rich microbe flora in the sperm population (‘sperm-associated microbiota’). These sperm-associated microbiota differed between the two species of cricket and between different populations and showed functional diversity. Further, sperm-associated microbiota killed sperm, highlighting their potential role in fitness, especially since they are most likely transferred to females during mating.

## Introduction

While studying aspects of the sperm biology and immunity of cricket species using the *in vitro* ejaculation technique (Figure 1A) combined with microbiological standard techniques, we encountered bacteria that were released from the male sperm container, the spermatophore, alongside sperm (Figure 1B). After extracting the spermatophore from the male, sterilizing its outside by drawing it through a flame or by ethanol, we cultivated microbes on agar plates and identified them (Supplementary Material). Examining these microbes in further detail, we discovered that i) microbes are regularly present in the spermatophore of crickets. We propose that this warrants the characterization as sperm-associated microbiota (SAM). ii) The current data suggest rich and diverse SAM. With six different and independent observation trials involving nine cricket populations (Table 1) and several protocols, we were able to characterize 65 morphotypes based on the external morphology of microbial colonies. It is important to note here that the same bacterial species may produce slightly different morphotypes across trials, protocols or other conditions. As expected, we isolated more unique 16S sequences from these 65 morphotypes, a total of 98 (Table 1). In addition to taxonomic diversity, the SAM were also functionally diverse. For example, we found differences in DNAse, lactase, lipase and protease activity (K. Reinhardt, unpublished data). iii) Importantly, using the sperm viability kit, we show that the microbe community isolated from the sperm container increased sperm mortality in the species or population they were sampled from (Fig 1) although as yet we are unable to assign the effect to any taxonomic or functional group.

**Figure 1.**
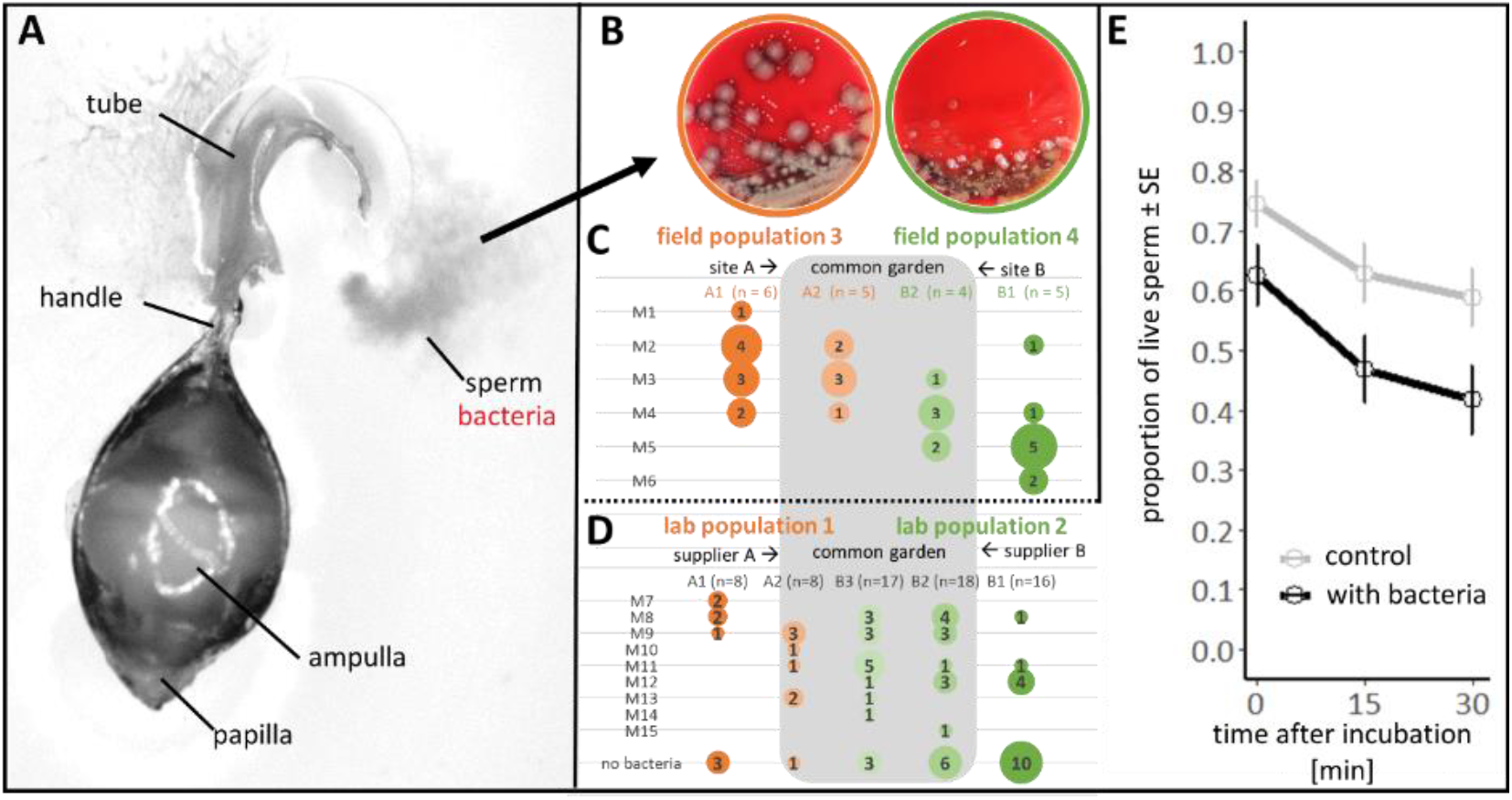
The sperm microbiome of crickets. Spermatophores (sperm containers) of male crickets can be used for in-vitro ejaculation on a microscope slide (A), showing a separation between sperm that leaves first, and the seminal fluid. The ejaculate contains microbes which differ between the cricket species and between different populations within species (B, field population 3 and 4). The bacteria are, at least partly of environmental origin: males of two populations, brought to the laboratory and sampled repeatedly, display increasingly similar sperm microbiome (M1-M15 represent different colony morphotypes, the circles and numbers represent the number of individuals in which this morphotypes was found, A1/B1 represents the first spermatophore, A2/B2 the second, and B3 the third spermatophore taken from a male?) in *G. campestris* (C) and *G. bimaculatus* (D). Bacteria, cultivated from the ejaculate of *G. bimaculatus* laboratory populations (lab population 1 and 2), increased sperm mortality in both laboratory populations (n=28-30) in comparison to sperm in a sterile control solution (E) (see supplement for method).

**Table 1.** List of bacteria detected in the SAM of two cricket species (*Gryllus* spec.). Bacterial identity is taken from 16S sequencing of bacteria (see supplement).

| cricket species | population | bacteria order | bacteria family | bacteria genus |
| --- | --- | --- | --- | --- |
| *G. bimaculatus* | commercial breeder 2 | *Lactobacillales* | *Aerococcaceae* | *Aerococcus* |
| *G. campestris* | field population 4 | *Lactobacillales* | *Aerococcaceae* | *Aerococcus* |
| *G. bimaculatus* | commercial breeder 1 | *Bacillales* | *Bacillaceae* | NA |
| *G. bimaculatus* | commercial breeder 2 | *Bacillales* | *Bacillaceae* | *Bacillus* |
| *G. bimaculatus* | lab population 1 | *Bacillales* | *Bacillaceae* | *Bacillus* |
| *G. campestris* | field population 3 | *Bacillales* | *Bacillaceae* | *Bacillus* |
| *G. campestris* | field population 4 | *Bacillales* | *Bacillaceae* | *Bacillus* |
| *G. campestris* | field population 4 | *Hyphomicrobiales* | *Brucellaceae/Phyllobacteriaceae* | NA |
| *G. bimaculatus* | lab population 1 | *Actinomycetales* | *Dermabacteraceae* | *Brachybacterium* |
| *G. bimaculatus* | lab population 1 | *Enterobacterales* | *Enterobacteriaceae* | *Enterobacter* |
| *G. bimaculatus* | lab population 1 | *Enterobacterales* | *Enterobacteriaceae* | *Enterococcus* |
| *G. bimaculatus* | lab population 1 | *Enterobacterales* | *Enterobacteriaceae* | *Klebsiella* |
| *G. bimaculatus* | lab population 2 | *Enterobacterales* | *Enterobacteriaceae* | *Enterococcus* |
| *G. bimaculatus* | lab population 3 | *Enterobacterales* | *Enterobacteriaceae* | *Enterococcus* |
| *G. campestris* | field population 3 | *Enterobacterales* | *Enterobacteriaceae* | *Enterococcus* |
| *G. campestris* | field population 4 | *Enterobacterales* | *Enterobacteriaceae* | *Enterococcus* |
| *G. campestris* | field population 1 | *Enterobacterales* | *Enterobacteriaceae* | *Enterococcus* |
| *G. campestris* | field population 1 | *Enterobacterales* | *Enterobacteriaceae* | *Klebsiella* |
| *G. campestris* | field population 2 | *Enterobacterales* | *Enterobacteriaceae* | *Klebsiella* |
| *G. bimaculatus* | commercial breeder 1 | *Pseudomonadales* | *Moraxellaceae* | NA |
| *G. bimaculatus* | lab population 3 | *Pseudomonadales* | *Moraxellaceae* | *Acinetobacter* |
| *G. campestris* | field population 4 | *Rhodospirillales* | *Rhodospirillaceae* | NA |
| *G. bimaculatus* | lab population 1 | *Bacillales* | *Staphylococcaceae* | *Staphylococcus* |
| *G. bimaculatus* | lab population 1 | *Bacillales* | *Staphylococcaceae* | *Staphylococcus* |
| *G. bimaculatus* | lab population 2 | *Bacillales* | *Staphylococcaceae* | NA |
| *G. bimaculatus* | lab population 3 | *Bacillales* | *Staphylococcaceae* | *Staphylococcus* |
| *G. campestris* | field population 3 | *Lactobacillales* | *Streptococcaceae* | *Lactococcus* |
| *G. campestris* | field population 4 | *Lactobacillales* | *Streptococcaceae* | *Lactococcus* |
| *G. campetris* | field population 1 | *Lactobacillales* | *Streptococcaceae* | *Lactococcus* |
| *G. campetris* | field population 2 | *Lactobacillales* | *Streptococcaceae* | *Streptococcaceae* |
| *G. bimaculatus* | lab population 1 | *Enterobacterales* | *Yersiniaceae* | *Serratia* |

iv) The SAM of crickets vary between populations, both in the field and the laboratory. One species, *Gryllus campestris*, displayed differences between two field sites near Tübingen (Germany) (ca. 4.5 km apart) and between two field sites near Dresden (Germany) (ca. 30 km apart). *Gryllus bimaculatus*, when obtained from different populations of commercial laboratory animal suppliers, also showed differences in their SAM (Table 1/ Fig. 1). We then revealed that at least some bacteria in the SAM are of environmental origin: Males brought into the laboratory and kept there for repeated sampling, showed a tendency in both abundance and species composition to converge towards highly similar SAM, and this notion was true for the two cricket species we investigated (Fig 1). Our attempts to clarify the infection route were not successful so far.

v) Within populations, males varied in the incidence, abundance and composition of the SAM (Table 1/ Fig 1). For example, in 56% of males (35 out of 63) we did not succeed in extracting cultivable bacteria. However, currently it is not clear to what extent this variation in incidence is related to host genetic variation or to environmental variation (such as the laboratory (see iii) above, Fig. 1D, E). We have not been able to systematically compare SAM in the spermatophore with that found in the sperm storage organs of females. However, our observations support the view that the existence of SAM may be the rule rather than the exception (Altmäe et al. 2019; Rowe et al. 2020). The high incidence of, and sperm mortality (Fig. 1E) caused by, sperm microbes that we discovered in our series of natural history observations suggest that the SAM is likely a significant selection factor that make the broad ecological significance put forward by these reviews plausible. Here, we briefly discuss how our observations may provide additional interpretations of biological phenomena in four areas of ecology and evolution - sexual selection, local adaptation, reproductive isolation, and ecological immunology.

Ecological immunology has shown that systemic immune responses or infection affect reproduction negatively (Table 2; Schwenke et al. 2016)) and, *vice versa*, that mating and reproduction influence immunity. However, the immunological consequences of mating and reproduction are not consistent across species, and neither are they across the types of immune responses (Schwenke et al. 2016, Oku et al. 2019). Moreover, studies frequently mention that effect of possible but unknown sexually transmitted microbes may contribute to mating-induced variation in immunity (Shoemaker et al. 2006; Schwenke et al. 2016). If SAM are as common in other animals as they are in crickets, they might be those long-sought after sexually transmitted microbes. Moreover, if SAM occur regularly, the idea of trade-offs between reproduction and immunity may need additional discussion because the management of regular sexually transmitted microbes requires a simultaneous investment in reproduction and immunity (Barribeau and Otti 2020).

**Table 2.** Selection of possible additional or alternative interpretations of cricket reproductive behaviour, immunity and sexual selection in the light of the existence of sperm-associated microbiota.

| Observation | Current interpretations | Possible alternative or additional interpretations |
| --- | --- | --- |
| Removal of the spermatophore by females is related to male quality (Zuk and Simmons 1997) | Cryptic female choice for male quality based on male or sperm quality | Females prevent the immigration of sperm-associated microbes |
| Mating changes female immune response (Oku et al. 2019, Schwenke et al. 2016) | (1) Immune suppression protects sperm from the female immune system; trade-off between immunity and reproduction.<br>(2) Immune activation viewed as a reduction of disease transmission risk. | More specifically: Immune activation protects females from infection by sperm-associated microbes. |
| Mating enhances parasite resistance (Shoemaker et al. 2006) | (1) females receive male compounds that enhance disease resistance (i.e. direct benefits)<br>(2) mating cues females to up-regulate immunity to respond to disease transmission | The female immune system has been upregulated to fight sperm-associated microbes. |
| Seminal fluid composition is very diverse (Simmons et al. 2013), contains prostaglandin synthetase (Larson et al. 2012) | Increased prostaglandin synthesis in females increases fertilization or egg-laying rates beyond female optimum/ females evolve resistance/ new male substances are selected | Increased prostaglandin synthesis in females increases antibiotic sperm protection from sexually transmitted microbes |
| Females frequently mate with the same male (Rodríguez-Muñoz et al. 2010) | Females receive fresh sperm (Reinhardt 2007) | Females may reduce the between-male diversity of sexually transmitted microbes |
| Many females in the wild leave no offspring (Rodríguez-Muñoz et al. 2010) | none | Microbe-induced sperm mortality affects fecundity |

Female investment into immunity likely depends on the composition of the SAM and the probability of sexual transmission. It is, therefore, possible that SAM assist in explaining differences between populations and species in the immunological consequences of mating. We believe it is a large, unexplored area to examine whether female defense against sexually transmitted microbes occurs by immune upregulation in anticipation to mating (Siva-Jothy et al. 2019) or by constitutive immune effectors. Future work in ecological immunology may, therefore, benefit from measuring immune responses just before as well as after mating.

The reproductive biology of crickets has long been a model system of behavioral ecology, condition-dependence and sexual selection research (Zuk and Simmons 1997). Briefly, as soon as the spermatophore is formed and deposited at a special pouch at the end of the abdomen, males start acoustic signaling to attract females. For mating, females place themselves over the male and from underneath the male attaches the spermatophore to the genital opening of the females. A long rod protruding from the spermatophore (Fig. 1A) is inserted into the female genital opening. Once the spermatophore is attached, sperm pumps out and travels into the female reproductive tract and sperm storage organ. While this is happening the male physically separates from the female and stays close by the female, a behavior known as guarding. Across several species of crickets, mate guarding delays the time at which the female removes the spermatophore and so increases the number of sperm migrating into the female. Several sexual-selection related hypotheses have been proposed for both the spermatophore removal by the female, as well as the guarding duration by the male (Zuk and Simmons 1997) (Table 2). However, necessarily all of them were tested in the absence of the knowledge of the existence of SAM reported here. Given that microbes associated with sperm cause sperm mortality (Fig. 1F), our observation may pave the way to the testing of additional hypotheses, some of which are proposed in Table 2.

We observed for both cricket species that microbes increased sperm mortality, that the SAM have an environmental component, and that SAM also vary within host species. The three observations together may allow for the hypothesis that males adapt to the sperm mortality imposed by the local microbes. If so, this would represent a form of local adaptation and represent a significant step towards any microbially-induced reproductive isolation proposed by Rowe et al. (2020). If such male adaptation would include the adaptation of ejaculate components (Otti et al. 2013), crickets and their SAM would represent a promising model of a possible coevolution between seminal fluid components and locally adapted, sexually transmitted microbes. This idea could have an influence on the interpretation of paternity outcomes in sperm competition trials involving males from the female’s own or separate population and may also help to explain conspecific sperm precedence in crickets (Gregory and Howard 1994), and perhaps other species. Because the SAM had an environmental component, the idea of sperm or ejaculate adaptation to SAM would allow for the prediction that ecologically more similar habitats harbor more similar sperm microbe communities. Therefore, SAM might even contribute to ecological speciation (Nosil 2012). Given the microbiota changes in the laboratory (Fig 1 D, E), our observations are, finally, a plea for field work.

## Supplementary material

### Isolation of microbes

The spermatophores were removed from the male genital opening using forceps. They were sterilised on the outside surface with ethanol or by flame treatment, placed in sterile medium using sterile forceps and crushed (Supplementary Table 1). After incubation, the medium was spread on sterile agar plates. Growing colonies were described morphologically and single bacterial strains were isolated and collected for 16S Sanger sequencing. DNA was isolated from these strains according to the respective protocol of the kit (see Supplementary Table 1), PCR were conducted to amplify variable regions of the 16 S DNA (see text below for details) and the PCR products were sent to a company for sequencing (see Supplementary Table 1). We used BLAST to determine the sequence identity of our bacteria samples in the NCBI database and assigned genus (94.5% sequence identity), family (86.5% sequence identity) and order (82.0% sequence identity) (Yarza et al. 2014). Note that only cultivable and (facultative) aerobic bacteria species can be detected by this method.

**Supplementary Table 1.**
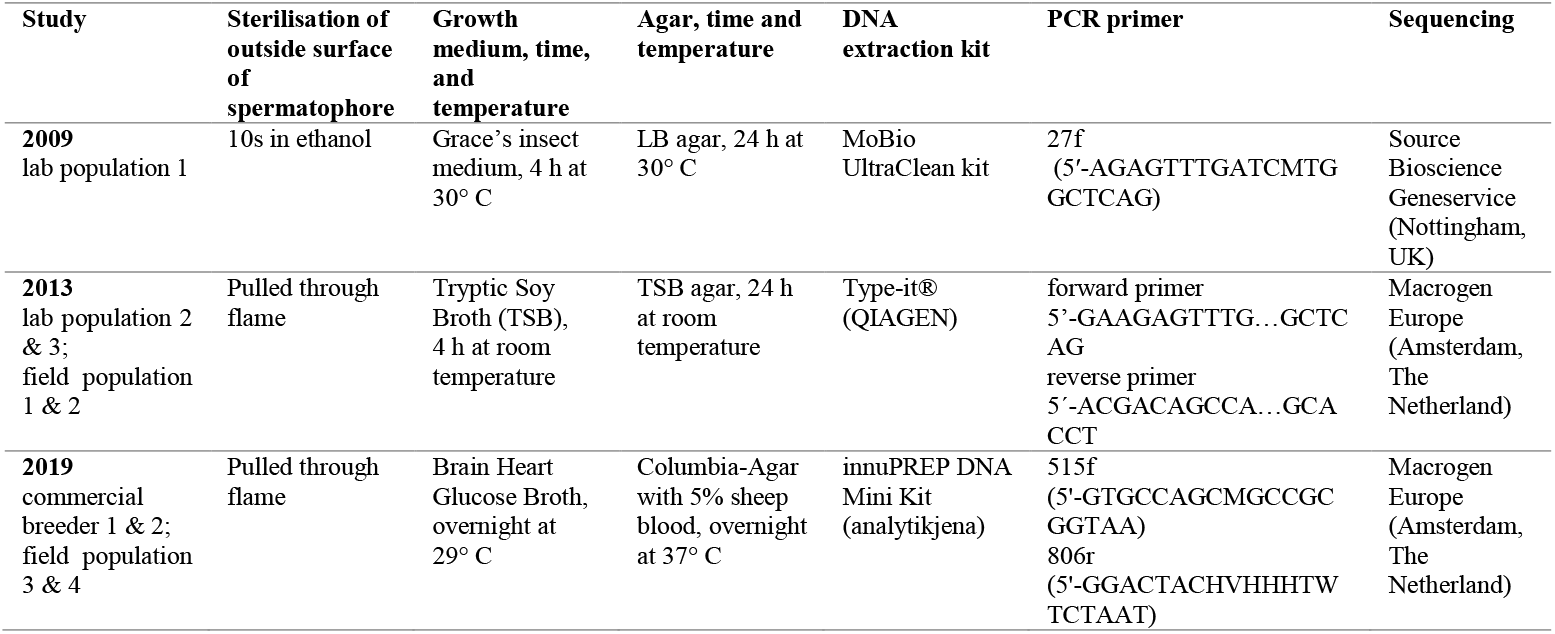
– Details of the different protocols used for the isolation of sperm-associated bacteria.

### Sperm viability assay

#### Bacterial solution

We made a bacterial solution for two cricket populations from commercial suppliers. Each bacterial solution contained the SAM isolated from ten spermatophore.

Work was done with sterile instruments in a sterile environment. Males of each population were randomly chosen. Spermatophores were removed and sterilised on the outside by drawing them through a flame. To control for contamination from the outside of the spermatophores, they were each washed in 50 μl brain heart glucose broth (BHG) (Thermo Scientific) on a microscopic slide. Then, they were placed in a 1.5 ml Eppendorf tube with 500 μl brain heart glucose broth and crushed. 10 μl of the control were spread on a Columbia agar plate with 5% sheep blood. The solution with the spermatophores and the plates were incubated at 29°C overnight. Then, the bacterial solutions produced from ten spermatophores with no bacterial growth on the plates controlling for contamination were mixed.. The bacterial solutions were stored in the fridge to inhibit bacterial growth during the experimental period. To achieve comparable conditions for both populations, the bacterial solutions were adjusted to an optical density of 0.3 A at 600 nm. During the experimental period, the bacterial density was controlled and adjusted at two-hour intervals.

#### Sperm viability assay

We used each 15 males of two laboratory populations of *G. bimaculatus*. Two spermatophores of each male were sampled and assigned to the control treatment and the bacterial treatment in randomised order. Spermatophores were removed from males and placed in 7 μl BHG on a microscopic slide. The spermatophore was opened to initiate the release of sperm. After ten minutes of sperm release, we removed the spermatophore and added 1.5 μl SYBR^®^14 and 2 μl propidium iodide to stain live cells fluorescent green and dead cells fluorescent red. After two minutes of staining, we added either 10 μl sterile BHG (control) or 10 μl bacterial solution (bacterial treatment) and measure sperm viability over time as described in Eckel et al, 2017.

### Cricket housing

2009: The *G. bimaculatus* population was maintained in transparent plastic boxes at 26oC and a 14:9h light:dark cycle with 30 min dawn/dusk periods. They were fed rat chow (7% cereal (wheat, maize, barley, wheat-feed) 15% vegetable proteins (soya bean meal), 5% animal protein (fish meal) and 3% vitamins (major and trace) and amino acids) ad libitum. Water was provided in cotton-plugged vials.

2013: *G. bimaculatus* populations were housed in transparent plastic boxes and *G. campestris* males were housed in single transparent plastic containers with a normal dark:light cycle corresponding to a central European summer and at 26°C. They were fed dry cat food (30% protein, 10% fat, 6.5% ashes, 3.5% crude fibre, 1.2% calcium, 1% phosphor and 10 mg/kg copper, 65 mg/kg zinc, 2 mg/kg iodide, 0.2 mg/kg selenium, 1000 mg/kg taurine and 15000 IE/kg vitamin A, 1500 IE/kg vitamin D3 and 150 mg/kg vitamin E) and oat flakes (13.5 g proteins, 58.7 g carbohydrates, thereof 0.7 g sugar, 7.0 g fat, thereof 1.3 g saturated fatty acids, 10.0 g dietary fibre and <0.01 g natrium) ad libitum. Water was provided in cotton-plugged vials.

2019: *G. bimaculatus* populations were housed in transparent plastic boxes and *G. campestris* males were housed in single transparent plastic containers at 20°C-25°C and 24 h light. They were fed dry cat food (32% protein, 15% fat, 6.5% raw ash, 5% crude fibre, 8% moisture content) ad libitum. Water was provided as aqua pearls with vitamin D (Hobby Terraristik, 1733).

## Notes

### Competing Interest Statement

The authors have declared no competing interest.

## References

Altmäe, S., J. M. Franasiak, and R. Mändar. 2019. The seminal microbiome in health and disease. Nature Reviews Urology 16:703–721.

Barribeau, S. M., and O. Otti. 2020. Sexual Reproduction and Immunity. eLS:1–10.

Gregory, P. G., and D. J. Howard. 1994. A post-insemination barrier to fertilization isolates two closely related ground crickets. Evolution 48:705–710.

Larson, E. L., J. A. Andrés, and R. G. Harrison. 2012. Influence of the Male Ejaculate on Post-Mating Prezygotic Barriers in Field Crickets. PLoS ONE 7.

Nosil, P. 2012. Ecological speciation. Oxford University Press, New York.

Oku, K., T. A. R. Price, and N. Wedell. 2019. Does mating negatively affect female immune defences in insects? Animal Biology 69:117–136.

Otti, O., A. P. McTighe, and K. Reinhardt. 2013. In vitro antimicrobial sperm protection by an ejaculate-like substance. Functional Ecology 27:219–226.

Reinhardt, K. 2007. Evolutionary consequences of sperm cell aging. The Quarterly Review of Biology 82:375–393.

Rodríguez-Muñoz, R., A. Bretman, J. Slate, C. A. Walling, and T. Tregenza. 2010. Natural and sexual selection in a wild insect population. Science 328:1269–1272.

Rowe, M., L. Veerus, P. Trosvik, A. Buckling, and T. Pizzari. 2020. The Reproductive Microbiome: An Emerging Driver of Sexual Selection, Sexual Conflict, Mating Systems, and Reproductive Isolation. Trends in Ecology and Evolution 35:220–234.

Schwenke, R. A., B. P. Lazzaro, and M. F. Wolfner. 2016. Reproduction-Immunity Trade-Offs in Insects. Annual Review of Entomology 61:239–256.

Shoemaker, K. L., N. M. Parsons, and S. A. Adamo. 2006. Mating enhances parasite resistance in the cricket Gryllus texensis. Animal Behaviour 71:371–380.

Simmons, L. W., Y. F. Tan, and A. H. Millar. 2013. Sperm and seminal fluid proteomes of the field cricket Teleogryllus oceanicus: Identification of novel proteins transferred to females at mating. Insect Molecular Biology 22:115–130.

Siva-Jothy, M. T., W. Zhong, R. Naylor, L. Heaton, W. Hentley, and E. Harney. 2019. Female bed bugs (Cimex lectularius L) anticipate the immunological consequences of traumatic insemination via feeding cues. Proceedings of the National Academy of Sciences of the United States of America 116:14682–14687.

Zuk, M., and L. W. Simmons. 1997. Reproductive strategies of the crickets (Orthoptera: Gryllidae). The evolution of mating systems in insects and arachnids 89:109.

## References

Eckel, B. A., R. Guo, and K. Reinhardt. 2017. More pitfalls with sperm viability staining and a viability-based stress test to characterize sperm quality. Front. Ecol. Evol. 5:165. Frontiers.

Yarza, P., P. Yilmaz, E. Pruesse, F. O. Glöckner, W. Ludwig, K.-H. Schleifer, W. B. Whitman, J. Euzéby, R. Amann, and R. Rosselló-Móra. 2014. Uniting the classification of cultured and uncultured bacteria and archaea using 16S rRNA gene sequences. Nat. Rev. Microbiol. 12:635– 645.

